# Protein folding stabilities are a major determinant of oxidation rates for buried methionine residues

**DOI:** 10.1101/2021.12.20.473526

**Authors:** Ethan J. Walker, John Q. Bettinger, Kevin A. Welle, Jennifer R. Hryhorenko, Adrian M. Molina Vargas, Mitchell R. O’Connell, Sina Ghaemmaghami

## Abstract

The oxidation of protein-bound methionines to form methionine sulfoxides has a broad range of biological ramifications, making it important to delineate factors that influence methionine oxidation rates within a protein. This is especially important for biopharmaceuticals, where oxidation can lead to deactivation and degradation. Previously, neighboring residue effects and solvent accessibility (SA) have been shown to impact the susceptibility of methionine residues to oxidation. In this study, we provide proteome-wide evidence that oxidation rates of buried methionine residues are also strongly influenced by the thermodynamic folding stability of proteins. We surveyed the *E. coli* proteome using several proteomic methodologies and globally measured oxidation rates of methionines in the presence and absence of tertiary structure, as well as folding stabilities of methionine containing domains. The data indicate that buried methionines have a wide range of protection factors against oxidation which correlate strongly with folding stabilities. Concordantly, we show that in comparison to *E. coli*, the proteome of the thermophile *T. thermophilus* is significantly more stable and thus more resistant to methionine oxidation. These results indicate that oxidation rates of buried methionines from the native state of proteins can be used as a metric of folding stability. To demonstrate the utility of this correlation, we used native methionine oxidation rates to survey the folding stabilities of *E. coli* and *T. thermophilus* proteomes at various temperatures and suggest a model that relates the temperature dependence of the folding stabilities of these two species to their optimal growth temperatures.

## Introduction

Sidechains of methionine and cysteine residues contain sulfur atoms that are readily oxidizable by reactive oxygen species (ROS). While not as well-studied as cysteine oxidation, methionine oxidation is an important enzymatically reversible post-translational modification (PTM) that has been shown to play a role in a number of physiological processes and cellular pathways (1–5). Methionine oxidation converts a hydrophobic residue to a polar residue and as such can greatly alter the biochemical and structural properties of proteins (1,2,6–9). Methionine oxidation can destabilize proteins and deactivate enzymes and thus is often considered a form of oxidative damage (10–12). This effect has been particularly well-documented in the context of aging and neurological diseases where dysregulation of antioxidant pathways and loss of proteostasis can bring about elevated levels of methionine oxidation and other detrimental protein modifications (3,13,14). It has also been shown that methionine oxidation can exacerbate protein misfolding or aggregation and worsen the symptoms of age-related diseases (15).

More recent studies have indicated that methionine oxidation is not solely a form of spontaneous protein damage and in certain contexts can modify the normal biological function of proteins in a regulated manner (16–19). For example, the oxidation of specific methionine residues in actin is catalyzed by the MICAL family of oxidoreductases that regulate cytoskeletal remodeling (5,20). It has also been suggested that methionine oxidation can act as an antioxidant scavenger in cells (1,2,4). This theory posits that surface methionines can become reversibly oxidized by ROS to prevent more dangerous, irreversible oxidation elsewhere in the same protein or other macromolecules. Importantly, unlike most other oxidatively modified macromolecules, oxidized methionines can be enzymatically reduced by methionine sulfoxide reductases (MSRs) providing a regeneratable sink for removal of damaging ROS (21).

Analysis of methionine oxidation is also of great significance to commercial production of protein therapeutics where trace levels of peroxides during production or storage can cause degradation and loss of activity. Antibody based therapeutics can be especially sensitive to methionine oxidation, making it important to understand and prevent this modification (22,23). Because levels of methionine oxidation can change significantly in different environments, it is important to monitor oxidation levels during development or long-term stability studies of biopharmaceuticals (24,25).

It has been shown that, within the same protein, different methionines have widely different susceptibilities to oxidation. As just one example, among the ten methionines in α1-antitrypsin, five are prone to oxidation and five are relatively protected from oxidation (26). The five oxidizable methionines in antitrypsin have widely different oxidation rates that are contingent on the conformation of the protein (27). Because of its importance to protein stability and function, it is important to understand the factors that influence the susceptibility of methionines to oxidation. There have been a number of published studies that have investigated factors that drive methionine oxidation (25,28–33). These studies have shown that solvent accessibility (SA) and neighboring sequence effects are two major determinants of methionine oxidation rates across the proteome. For example, a significant fraction of methionines are located near aromatic residues, which have been shown to reduce methionine oxidation rates *in vivo* (31,34). The fact that SA is a strong global predictor of methionine oxidation is supported by a number of studies showing that exposed methionines are typically oxidized faster than buried methionines (27,28,35,36). Indeed, this correlation has allowed methionine oxidation to be used as an experimental biochemical probe to monitor structural changes in proteins (37). However, a number of studies have suggested that SA and neighboring residue effects may not be sufficient to fully predict the propensity of methionines towards oxidation. Specifically, it has been argued that the oxidation of buried methionines may also be influenced by the dynamics of the native structure and conformational flexibility (29).

In this study, we considered the potential role of thermodynamic folding stabilities in determining the oxidation rates of buried methionines. Within a cell, most protein domains exist in a dynamic equilibrium between folded and unfolded conformations (38–40). The free energy difference between these two states (ΔG_folding_ or ‘thermodynamic folding stability’) establishes the fraction of its population that is in a folded conformation at equilibrium (41). For some buried methionines, transient protein unfolding may be necessary to expose the protein to oxidation. Thus, folding stability can modulate the folding equilibrium and establish the rate of methionine oxidation for buried methionines. To date, correlations between oxidation rates and protein stabilities have been reported for a few individual proteins (42,43). However, the general correlation between folding stability and methionine oxidation rates has not been studied on a proteome-wide scale.

We used MS-based proteomics to quantify oxidation rates of methionines within folded and unfolded proteins and calculated relative oxidation protection factors (PFs) for methionines on a global scale. We then measured folding stabilities for the detected proteins using SPROX, a proteome-wide methodology for conducting denaturation experiments (37,44). The resulting largescale datasets allowed us to explore the correlation between PFs and folding stabilities within the proteomes of the mesophile *E. coli* and the thermophile *T. thermophilus*. The analysis globally quantified the influence of folding stabilities on methionine oxidation rates and provided insights into the relationship between proteome stabilities and optimal growth temperatures of these two bacterial organisms.

## Experimental Procedures

### Culture growth and lysate preparations

*Escherichia coli* K-12 W3110 was a generous gift from Dr. Gloria Culver (University of Rochester), and was cultured in lab-made lysogeny broth at 37°C. *Thermus thermophilus* strain HB8 (ATCC) was cultured at 70°C in Thermus medium (ATCC medium 697) made in house. For both species, initial overnight growths were streaked onto agar plates, from which single colonies were picked for 200-mL batches, grown to late-log phase, and aliquoted into glycerol stocks at −80°C. All further cultures were propagated from these stocks. For both species, 5- or 10-mL overnight cultures were diluted at a ratio of 1:100, grown to late-log phase, and collected on ice. Resulting cell pellets were immediately frozen at −80°C until use. All cells were lysed at 4°C using gentle sonication in a native lysis buffer (EDTA-free protease inhibitor mini tablets (Pierce) in filtered 20 mM sodium phosphate buffer (pH 7.4) with 50 m*M* NaCl), with resting periods on ice. Protein concentration was subsequently measured using a bicinchoninic acid (BCA) assay and standardized to a final concentration of 2.5 mg/mL for all intact protein experiments. After standardization, lysates were aliquoted and flash-frozen in liquid N2 and kept at −80°C until use.

### Peptide oxidation

Proteomic peptide samples were prepared from native lysate, as above, which was immediately purified and digested with trypsin (see *Sample purification and TMT labeling* section below). Oxidation was then performed on the purified peptides in a manner similar to that published previously (45). Peptides were brought to a final concentration of 1 mg/mL, and oxidative labeling was accomplished using a concentration-gradient of heavy-labeled H_2_^18^O_2_ (Cambridge Isotope Laboratories). Specifically, samples were treated with 0, 0.02, 0.04, 0.08 or 0.16 *M* O-18 peroxide, for 30 min at 37°C in a heat block. All samples were quenched using an excess of 1 *M* sodium sulfite and lyophilized. Quenching reagent was removed via desalting using lab-made C18 spin columns. Columns were first conditioned using elution buffer containing 50% acetonitrile (ACN) in 0.1% trifluoroacetic acid (TFA) (Thermo Scientific) and washed with sample buffer (0.1% TFA in water). Samples were resuspended in sample buffer, added to the columns, and washed. After elution, samples were lyophilized, then oxidized using a final concentration of 0.16 *M* H_2_^18^O_2_ for 30 min at 37°C to block any remaining methionines. Samples were lyophilized again to remove peroxide and resuspended in sample buffer for MS analysis.

### Native oxidation

Samples were thawed on ice, then incubated for 10 min at RT with a final concentration of 100 μ*M* sodium azide to prevent the degradation of peroxide by native catalases. Oxidation was achieved via a time course, as this was more amenable to measuring the slow rates expected for buried residues. Time courses were optimized for their given temperatures to account for the change in kinetics, and exact time points are listed in Supp. Table 1. For each set of samples, a native, unoxidized control and a fully denatured, oxidized (final concentration of 5% sodium dodecyl sulfate (SDS) heated to 90°C for 10 min) control were also prepared. Samples were oxidized using a final concentration of 0.98 *M* peroxide for 25°C and 37°C, or 0.245 *M* for 50°C, then purified as described in below in *Sample purification and TMT labeling*. An Amplex Red (Invitrogen) hydrogen peroxide assay kit was used to validate the steady state levels of peroxide in samples treated with 100 μ*M* sodium azide under each condition (data not shown). To capture the large range of rates expected for these samples, it was advantageous to adopt a labeling method more amenable to high-throughput analysis. Thus, we opted for using Tandem mass tag (TMT) 10-plex (see TMT labeling section below) reagents to quantify the fraction of oxidation in multiplexed samples. This allowed us to maximize coverage by using peptide fractionation protocols while cutting down on the number of MS runs.

### SPROX

SPROX oxidation, denaturation, and sample cleanup were performed as described previously (44). The guanidinium chloride (GdmCl) concentrations used for each experiment are listed in Supp. Table 1. Stability measurements was performed using quality control filters as described previously (44).

### Sample purification and TMT labeling

Native oxidation and peptide samples were purified using an S-Trap mini kit 2.1 (ProtiFi) using manufacturer’s protocol. 100 m*M* iodoacetamide was used as the alkylating agent and overnight trypsin digests were conducted at a trypsin-protein ratio of 1:25 at 37°C. SPROX sample cleanup was performed as previously described (44) and digested overnight with a trypsin-protein ratio of 1:100 at 37°C. After digestion, TMT labeling for SPROX and native oxidation samples was conducted with a TMT10plex mass tag labeling kit (Thermo-Scientific), using 0.2 mg of isobaric label per 10 μg of lysate digest.

### Peptide fractionation by spin column

To increase coverage, all peptide samples were fractionated using C18 spin columns prior to LC-MS/MS analysis. Columns were first conditioned using ACN and 100 mM ammonium hydroxide (AH). Samples were resuspended in 50-100 μL of AH and added to the columns. After washing the columns with ACN and AH, samples were eluted with a gradient of 16 concentrations of ACN ranging from 2-50%. Eluent fractions were then recombined into 8 fractions in a staggered manner (#1 with #8, #2 with #9, etc.), dried down and resuspended in 0.1% TFA at a concentration of 0.25 mg/ml for LC-MS/MS analysis.

### LC-MS/MS

TMT-labeled peptides originating from the native oxidation and SPROX methods were injected onto an Easy nLC-1200 HPLC instrument (Thermo Fisher) using a 30 cm C18 column packed with 1.8 μm beads (Sepax), made in-house, for analysis via a Fusion Lumos Tribrid mass spectrometer (Thermo Fisher). Peptides were eluted from the column using a gradient of two solvents: A (0.1% formic acid in water) and B (0.1% formic acid in 80% ACN). The gradient started with 3% B for 2 min, increased to 10% B over 7 min, then to 38% B over 94 min, finally ramped up to 90% B over 5 min, and held for 3 min, before returning to starting conditions in 2 min and re-equilibrating for 7 min, for a total run time of 120 min. Ionization was carried out using a Nanospray Flex source operating at 2 kV. The Fusion Lumos was operated in data-dependent mode, acquiring both MS1 and MS2 scans in the Orbitrap with a cycle time of 3 s. Monoisotopic Precursor Selection (MIPS) was set to Peptide. The scan range was 400-1500 *m/z*, with an AGC target of 4e5, a resolution of 120,000 at *m/z* of 200, and a maximum injection time of 50 ms. Only peptides with a charge state between 2-5 were picked for fragmentation. Precursor ions were fragmented by higher-energy collisional dissociation (HCD) using a collision energy of 38% and an isolation width of 1.0 *m/z*. MS2 scans were collected with a resolution of 50,000, a maximum injection time of 105 ms, and an AGC setting of 1e5. Dynamic exclusion was set to 45 s.

Analysis of samples generated by the peptide oxidation experiment was conducted as above with the following changes. The runtime was reduced to 90 minutes, while the cycle time was set to 1 second to ensure enough scans across the peak for accurate MS1 quantitation. Precursor ions were fragmented by collision-induced dissociation (CID) using a collision energy of 30 and an isolation width of 1.1 *m/z*. MS2 scans were collected in the ion trap with the scan rate set to rapid, a maximum injection time of 35 ms, and an AGC setting of 1e4. Dynamic exclusion was set to 20 s.

### Database searches and quantitation

For SPROX and native oxidation experiments, MS2 data were searched against the *E. coli* (4,352 entries, downloaded 1/18/2019) or *T. thermophilus* (2,227 entries, downloaded 1/18/2019) UniProt databases using the integrated Andromeda search engine with MaxQuant software (46). Searches included the TMT10-plex labels as fixed modifications and methionine oxidation as variable modifications and allowed for up to two missed cleavages with maximum false discovery rate (FDR) thresholds of 1%. Resulting ion intensities were then normalized using total channel intensity of non-Met peptides and median normalization. MS searching, quantification and curve fitting to acquire intrinsic rate constants for proteomic *E. coli* peptides was performed as previously described (45). Coverage for all MS experiments is shown in Supp. Table 2.

### Solvent accessibility (SA) measurements

SA measurements from Protein Data Bank (PDB) structures were computed for available *E. coli* and *T. thermophilus* proteins as previously described (44). Briefly, solvent accessible surface areas of methionine residues were calculated in Pymol, with dot density set to three. If a protein was represented by more than one PDB file, the value reported is the median value for all structures. A maximum of three PDB files were analyzed for each protein. Multimeric structures were analyzed with all subunits present. Additionally, SA values for *E. coli* were computed using structures available from the AlphaFold (AF) Protein Structure Database using AlphaFold2 (47). Structures for *T. thermophilus* were not available in the AF database. Therefore, they were predicted using ColabFold (48), utilizing the fast homology search of MMseqs2 with AlphaFold2, with the AlphaFold2_advanced option set to the default parameters for all proteins. This analysis was conducted using the computational resources of the University of Rochester’s Center for Integrated Research Computing (CIRC) Linux cluster. The top five ranked models for each protein were then used to calculate SAs as above, keeping only models with an average plDDT value >70. Where SA measurements for both PDB and AF structures were available, the two values were averaged. SA measurements are reported in Supp. Table 3.

### Measurements of oxidation rate constants and protection factors

Pseudo-first order observed rate constants for oxidation of peptides and proteins (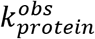 and 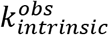, respectively) were measured by least-squares fitting of fractional oxidation (*f_ox_*) of each methionine containing peptide to the following equation:

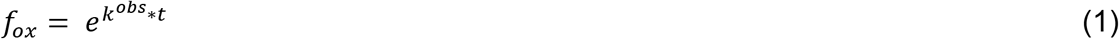

Where *t* is the oxidation time in minutes. Given that concentration of H_2_O_2_ was in excess of methionines in all experiments, the above pseudo-first order rate constants can be converted to second-order oxidation rate constants for proteins and peptides (*k_protein_* and *k_intrinsic_*, respectively) by dividing k^obs^ values by the molar concentration of H_2_O_2_. For intact proteins, protection factors (PFs) were measured by taking the ratio of *k_protein_* and *k_intrinsic_*:

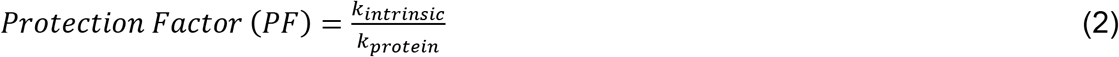

Since measured peptide *k_intrinsic_* values closely matched the oxidation rate of free methionine (23), we used the latter as our reference *k_intrinsic_* value for measuring PFs (after correcting for temperature using the Arrhenius equation (43)). For *E. coli*, where it was possible to calculate PFs for a portion of the data using *k_intrinsic_* values derived from peptide rate measurements, the measured PFs were very similar to those measured using *k_intrinsic_* values derived using free methionine. We chose to use the latter as it significantly increased the number of measured PFs (as there was limited overlap between proteomic coverages of peptide and protein oxidation experiments). Since our time course was optimized to survey slow, protected methionines, the maximum value of *k_protein_* in each experiment was set to the respective value for *k_intrinsic_*. Rates for proteins and peptides were kept only for tryptic peptides containing a single Met, which also lacked Cys, and had a *r*^2^ (goodness-of-fit to Eq. 1) ≥ 0.75. The filtered rate data and resulting PFs are listed in Supp. Table 3.

### Model for relating PFs to folding stabilities

For protected methionines where protein unfolding is required for oxidation, the oxidation reaction can be modeled based on the following reaction:

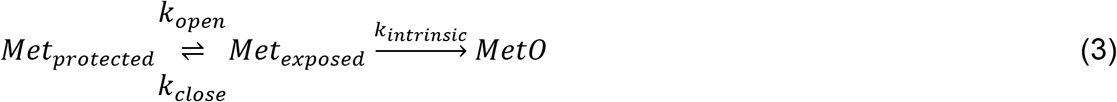

Where *k_open_* and *k_close_* are the folding and unfolding rate constants, respectively, and can be related to the folding equilibrium constant:

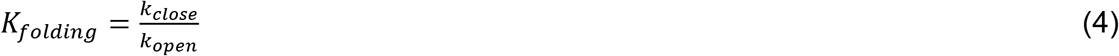

Analogous to the model used for analysis of hydrogen-deuterium exchange (HDX) experiments (49,50), the following relationship is established between the observed rate constant for oxidation (*k_protein_*) and the three rate constants employed in the model:

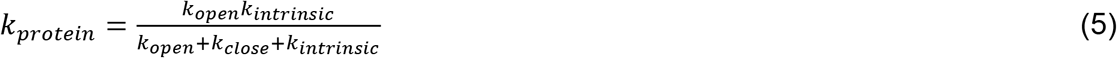

A survey of available protein folding rates from the protein folding database (PFDB) indicated that the average protein has a folding rate roughly 15,000 times faster than the oxidation of free methionine (51). Therefore, this reaction is expected to be in an EX2 regime (49,50) where *k_close_* is significantly faster than *k_open_* and *k_intrinsic_*, and therefore:

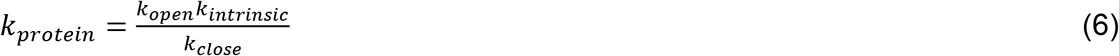

From equations 2, 4 and 6:

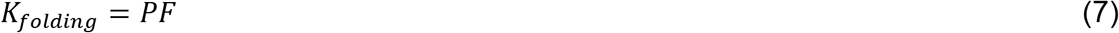

PFs can thus be related to folding stabilities (ΔG_folding_) and midpoints of denaturation curves (C_1/2_):

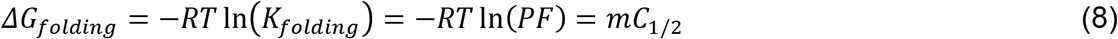

Where *R* is the gas constant, *T* is the temperature (in Kelvin) and *m* is the slope of the linear relationship between ΔG_folding_ and denaturant concentration in accordance to a two-state folding model (44). Therefore, if the assumptions of the above model are valid, oxidation PFs are expected to be correlated with folding stabilities and C_1/2_ values.

## Results

### Overview of experimental strategy

Oxidation of protein-bound methionines that are fully or partially solvent-exposed can potentially occur from the folded state of the protein without transient unfolding of the native structure. For these exposed methionines, SA and neighboring residue effects may strongly influence the rate of oxidation. However, we reasoned that for buried methionine residues, transient unfolding of the native structure is required to expose the sulfur atom to oxidation (Fig.1A). In this scenario, transient protein unfolding may be the limiting step in the oxidation process. Accordingly, under conditions where the folding reaction is in a rapid pre-equilibrium relative to the oxidation reaction, the folding equilibrium constant will be directly correlated to the rate of oxidation (Fig.1A and Discussion). This concept is analogous to the EX2 model commonly used to interpret HDX experiments where the rate of exchange of hydrogen-bonded backbone protons are known to correlate with folding stabilities (50). Thus, primary sequence, SA and folding stability can all potentially influence the rate of methionine oxidation, and the relative contribution of each parameter may depend on the conformational properties of the methionine sidechain. To quantify the contribution of each of these parameters to oxidation propensities of protein-bound methionines, we conducted three orthogonal proteomic experiments (Fig. 1B) to globally measure a) intrinsic rates of methionine oxidation within unstructured peptides, b) rates of methionine oxidation within intact native proteins, and c) folding stabilities of methionione-containing protein domains.

**Figure 1.**
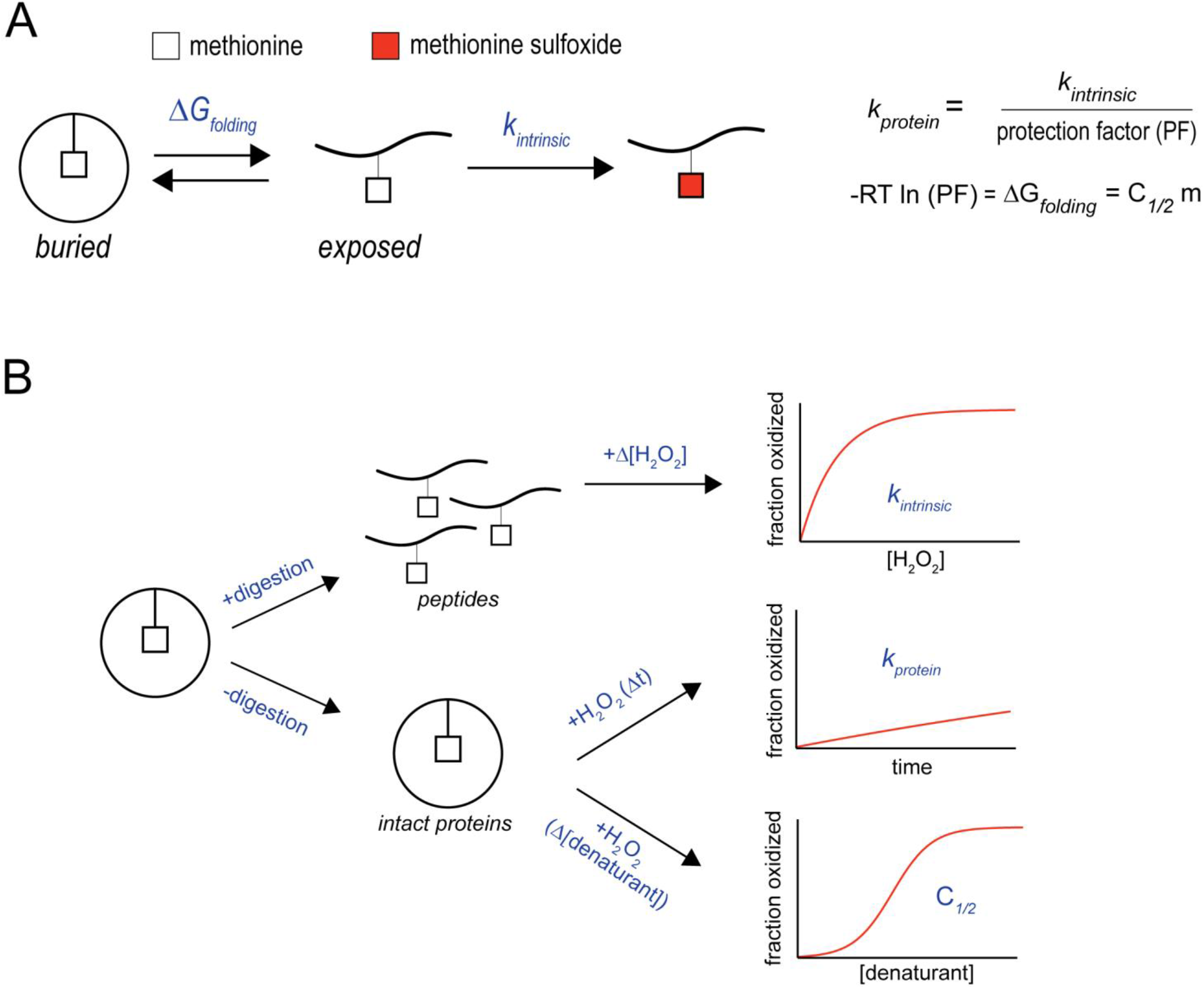
Schematic representations of proposed model and methods. A) For buried methionines, protein unfolding precedes oxidation. Oxidation protection factors (PFs) of buried methionines are defined as the ratio of methionine oxidation rates from the unfolded state (*k_intrinsic_*) and the observed oxidation rate (*k_protein_*). According to a two-state folding model, PFs are related to folding stability (ΔG_folding_) and the midpoint of denaturation (C_1/2_) as indicated in the right side of the figure. B) The three proteomic methods employed in this study. Intrinsic oxidation rates (*k_intrinsic_*) were measured using trypsin-digested proteins. Oxidation rates in intact proteins (*k_protein_*) were measured using native protein lysates. Folding stabilities (C_1/2_ values) were measured by SPROX. This method exposes native extracts to increasing concentrations chemical denaturants prior to oxidation.

### Intrinsic methionine oxidation rates within unstructured peptides

To measure intrinsic rate constants for methionine oxidation in unstructured peptides (*k_intrinstic_*), *E. coli* protein extracts were digested with trypsin prior to treatment with H_2_O_2_ (Fig. 2A). Peptides were treated with varying concentrations of ^18^O-labeled H_2_O_2_ at 37°C for 30 minutes and were subsequently blocked by the addition of excess levels of ^16^O-labeled H_2_O_2_. Fractional oxidation levels were determined with LC-MS/MS by quantifying relative intensities of ^18^O- and ^16^O-oxidized peptides using a bottom-up proteomics workflow. As previously described, this approach allows for accurate quantitation of methionine oxidation while minimizing the effects of spurious oxidation that could occur during sample processing and LC-MS/MS analyses (45). *k_intrinsic_* values were measured by analyzing the relationship between fractional oxidation and concentrations of ^18^O-labeled H_2_O_2_ treatments using a first-order kinetic model. We limited our analyses to peptides containing single methionines and no cysteine residues in order to minimize the confounding effects of multiple methionine oxidation events and cystine oxidation. In all, we were able to measure *k_intrinsic_* values for approximately 3,900 methionine-containing peptides (Fig. 2B). Measured *k_intrinsic_* values spanned approximately one order of magnitude with a median of 1.02 *M*^−1^ min^−1^, which is in line with previously measured oxidation rates for free methionine at this temperature (23,43).

**Figure 2.**
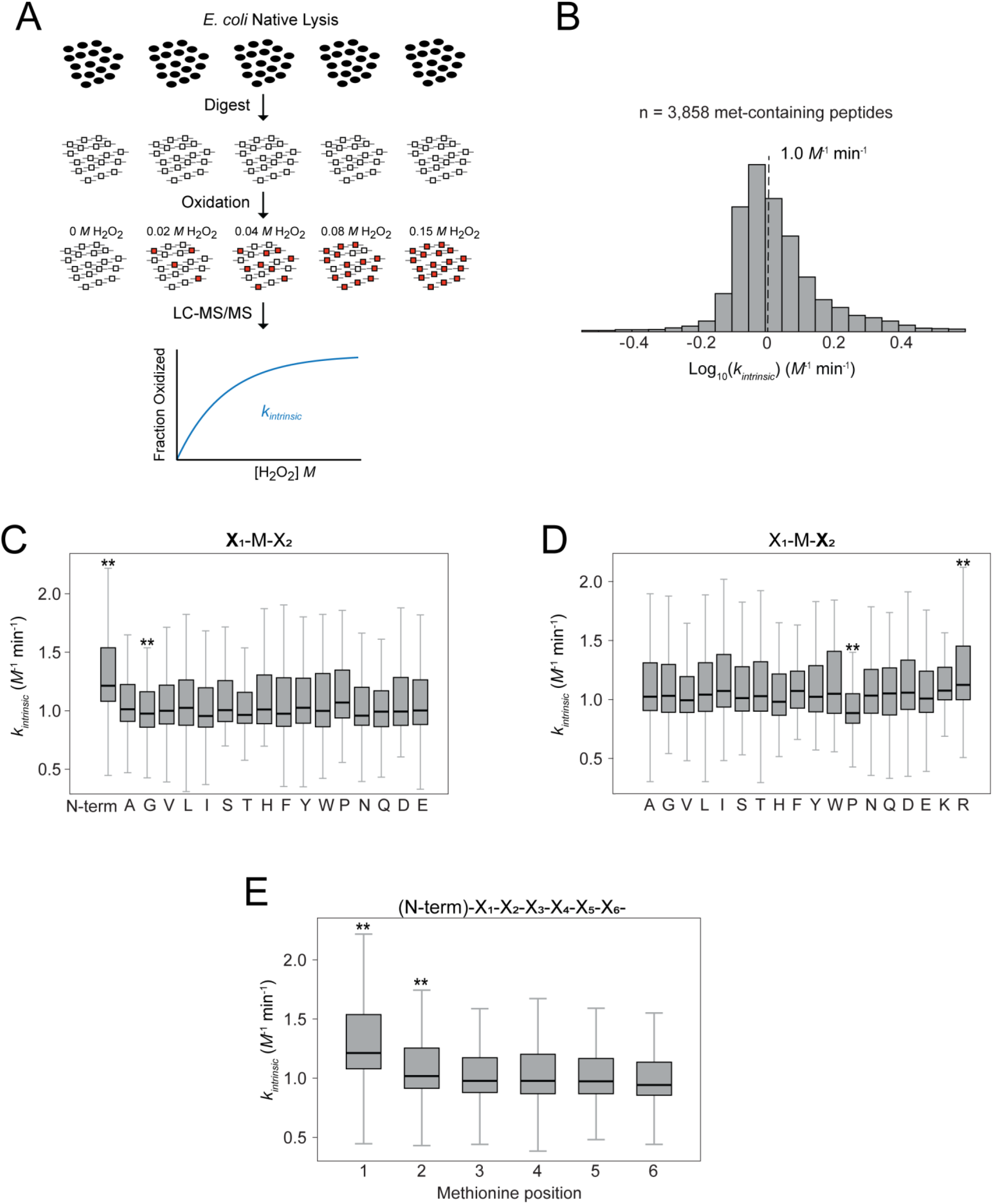
*k_intrinsic_* measurements and neighboring residue effects in the *E. coli* proteome. A) Experimental method. Trypsin-digested peptides samples were exposed to varying concentrations of H_2_O_2_. Fractional oxidation levels were measured with LC-MS/MS and used to calculate pseudo-first order oxidation rate constants. In the schematic, closed ovals indicate methionines in the native state, open squares indicate exposed methionines in peptides and red squares indicate oxidized methionines. B) Relative distribution of *k_intrinsic_* measurements in the *E. coli* proteome. The dotted line indicates the distribution median. C, D) Effect of neighboring residues on *k_intrinsic_* of methionines. E) Effect of position relative to the N-terminus on *k_intrinsic_* of methionines. For C-E, boxes indicate the interquartile range and whiskers show the entire range of values excluding outliers (> 2 SD). ** indicates a p-value < 0.001 using a Mann-Whitney U test, corrected for family-wise error rate with a Holm–Šidák method (significance level = 0.001).

We then used our proteome-wide data to analyze the effects of primary sequence on *k_intrinsic_* values (Fig. 2C-E). We observed that methionines located at the N-termini of peptides have significantly faster oxidation rates in comparison to methionines that are internal in the sequence (Fig. 2E). Other neighboring residue effects were comparatively less pronounced. For example, the presence of prolines at +1 positions and glycines at −1 positions of methionines correlated with slightly slower oxidation rates, whereas the presence of arginines at +1 positions correlated with slightly faster oxidation rates. It should be noted that because peptides were generated by trypsin digestion (which hydrolyzes proteins at the carboxyl side of lysines and arginines), certain neighboring residue combinations (e.g. arginines and lysines at −1 positions and methionines at C-termini) could not be observed in our analyses. Furthermore, methionines that are at −1 positions relative to lysines and arginines are necessarily positioned near the C-termini of peptides which may bias their oxidation rates. Nonetheless, our analysis indicates that regardless of primary sequence context, most protein-bound methionines in unstructured peptides are oxidized at relatively similar rates that are close to oxidation rates of free methionines.

### Rates of methionine oxidation within intact proteins

To investigate the effects of higher order structure on methionine oxidation, we next measured methionine oxidation rates within intact proteins (*k_protein_*) by exposing *E. coli* native extracts to H_2_O_2_ prior to digestion and LC-MS/MS analyses (Fig. 3A). To facilitate these experiments, two additional changes were made to the experimental protocols described above. First, oxidation reactions were carried out as a function of time rather than as a function of H_2_O_2_ concentration. Second, we took advantage of an isobaric tagging strategy (TMT) to multiplex the larger number of samples analyzed in this experiment (See Experimental Procedures for additional details and rationale for these protocol modifications).

**Figure 3.**
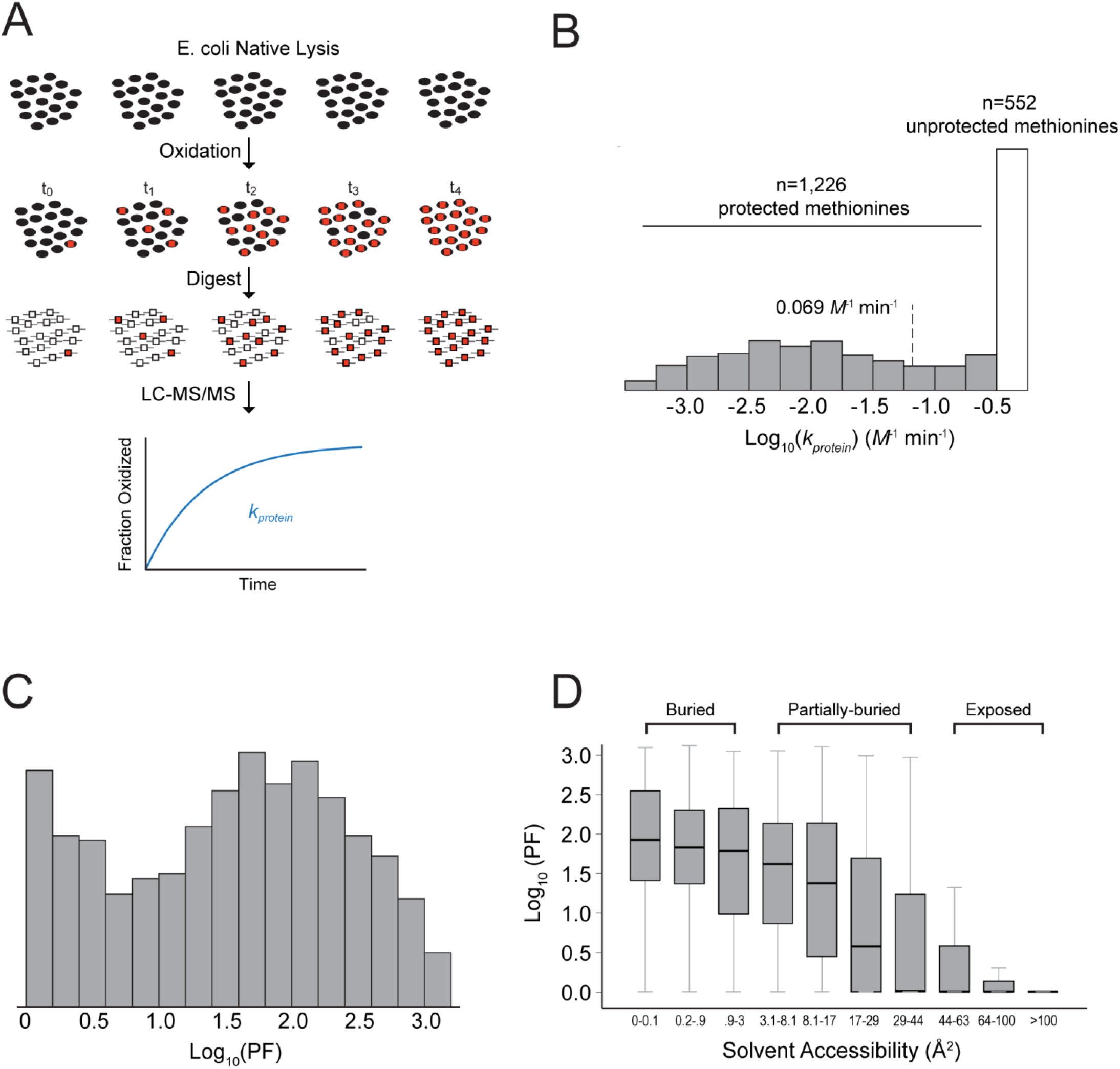
*k_protein_* measurements within native proteins and correlations with solvent accessibility (SA). A) Experimental method. Native lysates were treated with H_2_O_2_ using a range of oxidation times. Fractional oxidation levels were measured with LC-MS/MS and used to calculate pseudo-first order oxidation rate constants. In the schematic, ovals indicate methionines in the native state, squares indicate exposed methionines in peptides, black indicates unoxidized methionines and red indicates oxidized methionines. B) The relative log_10_ distribution of *k_protein_* measurements. The dotted line indicates the distribution median. Gray and white bars indicate protected and unprotected methionines as defined in the text. C) The relative distribution of PFs in the *E. coli* proteome. D) The relationship between PF and SA measurements. Methionines were categorized into three groups (buried, partially-buried and exposed) based in their SA values. Boxes indicate the interquartile range and whiskers show the entire range of values excluding outliers (> 2 SD). The total pairwise comparison between PFs and SAs had a significant level of correlation (Spearman rank test *ρ* <0.001).

We were able to measure native oxidation rates (*k_protein_*) for 1778 methionines mapped to 831 different proteins in the *E. coli* proteome. Among these methionines, ~600 were oxidized at fast rates that were within the range of peptide *k_intrinsic_* measurements and free methionines (Fig. 3B). We refer to these residues as “unprotected” methionines. The remaining ~1,200 “protected” methionines are oxidized at significantly slower rates in comparison to unstructured peptides. The overall range of *k_protein_* measurements spans at least 3 orders of magnitude, representing a significantly broader distribution than *k_intrinsic_* values. This observation indicates that for most protected methionines in proteins, higher order structure, and not primary sequence, is the dominant factor in establishing oxidation rates. The distribution of measured PFs for protected methionines (the ratio of *k_protein_* and *k_intrinsic_* measurements for individual methionines) is plotted in Figure 3C. According to the simplified two-state unfolding model illustrated in Figure 1A and described in Experimental Procedures, PF values are expected to be equal to the folding equilibrium constant and are logarithmically correlated with thermodynamic folding stabilities (ΔG_folding_).

We next examined the relationship between methionine SAs and PFs. SA values for methionines were calculated as described previously (44) using either their experimentally determined structure in the Protein Data Bank (PDB) or as predicted by Alphafold2 (47). In total, we were able to calculate both PF and SA values for 1731 distinct methionines in *E. coli*. Overall, there is a significant statistical correlation between PF and SA (Spearman Rank Correlation *ρ* <0.001). A closer examination of the relationship between SA and PF values indicated that methionine residues can be categorized into one of three general categories based on their oxidation properties (Fig. 3D). Highly exposed methionines (SA ≳ 44 Å^2^) are oxidized rapidly at rates that are equivalent to free methionines (PF ≈ 1). Partially buried methionine residues (SA ≈ 3 to 44 Å^2^) have a range of PF values that are correlated to their solvent accessibility. Buried methionine residues (SA ≲ 3 Å^2^) have high PF values that are independent of solvent accessibility.

The above data suggest that oxidation rates of fully exposed methionines are likely dictated by their primary sequence context. For partially-buried methionines, oxidation is correlated to SA established by the native structure of the protein. This suggests that for this class of methionines, oxidation likely occurs from the native state and does not require transient conformational unfolding of the protein structure. For fully-buried methionines, oxidation rates are significantly slower and independent of SA. Yet, these methionines have a wide range of PFs spanning three orders of magnitude. We reasoned that for this class of methionines, oxidation occurs from the unfolded state of the protein and hence transient protein unfolding must precede oxidation. Hence, we hypothesized that for fully-buried methionines, oxidation rates may be correlated to protein thermodynamic folding stabilities.

### Thermodynamic folding stabilities of methionine-containing protein domains

We used the proteomic methodology SPROX to conduct denaturation experiments on *E. coli* extracts and quantify protein folding stabilities on a global scale (Fig. 4A). In SPROX, proteins are gradually unfolded by the addition of increasing concentrations of a chemical denaturant such as GdmCl and subsequently pulse-oxidized by exposure to H_2_O_2_ (37,44). Measurements of relative fractional oxidation as a function of denaturant concentration are used to generate denaturation curves and determine C_1/2_ values (the denaturant concentration that induces the unfolding of half of the protein population). Protein domains with higher C_1/2_ values have higher conformational stabilities and lower ΔG_folding_ values (See Experimental Procedures for a detailed description and assumptions of the folding model).

**Figure 4.**
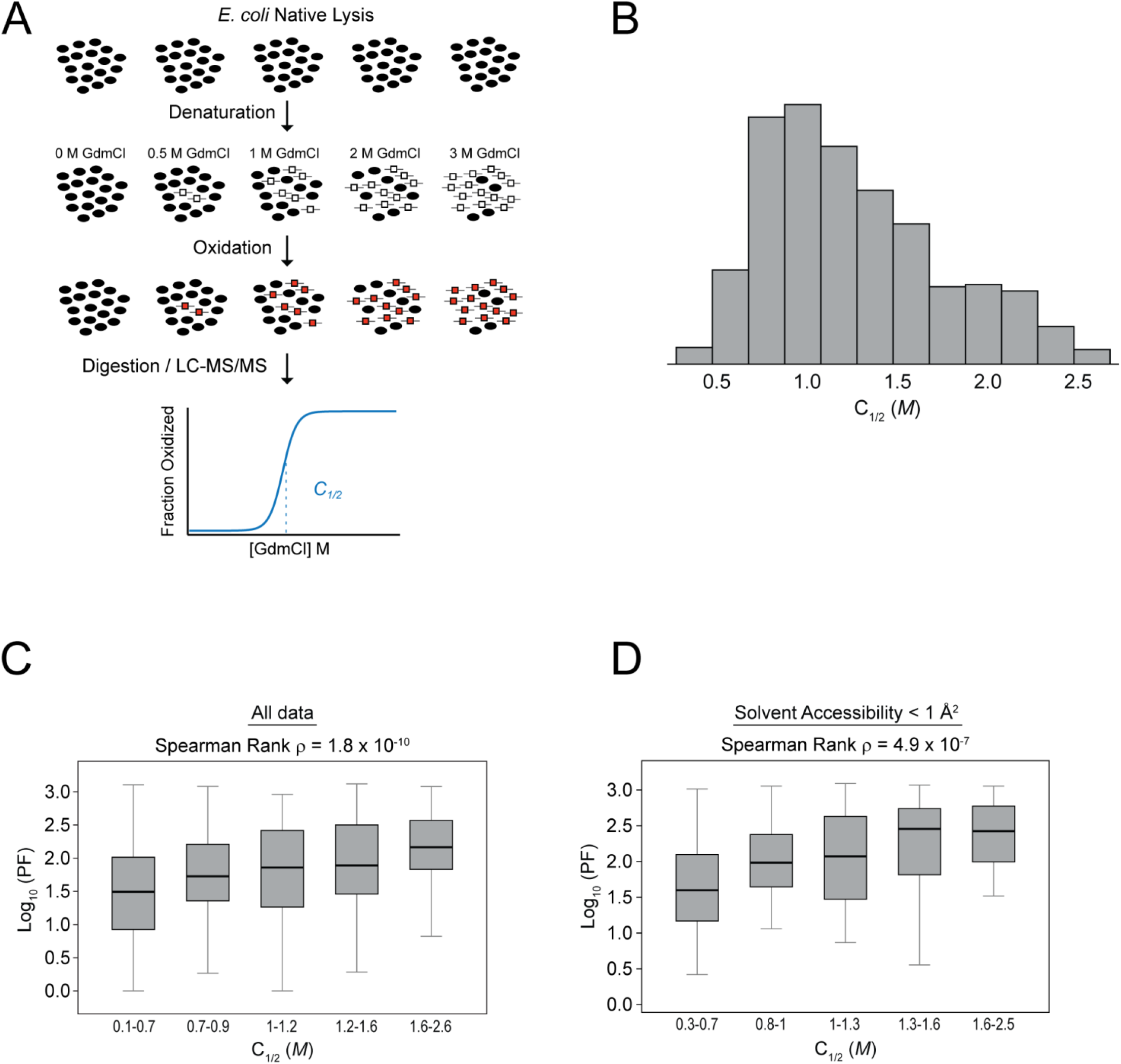
C_1/2_ measurements within native proteins and correlations with PFs. A) Experimental method (SPROX). Native lysates were treated with increasing concentrations of denaturant (GdmCl) prior to oxidation with H_2_O_2_ and subsequent digestion with trypsin. Fractional oxidation levels were measured with LC-MS/MS and used to measure C_1/2_ values. Symbols in the schematic are described in Figure 3. B) The relative distribution of C_1/2_ values in the *E. coli* proteome. C, D) The relationship between PF and C_1/2_ values for all methionines (C) and highly protected methionines (D). Boxes indicate the interquartile range and whiskers show the entire range of values excluding outliers (> 2 SD). The Spearman rank correlation values indicate the correlation for total pairwise comparisons.

Using this method, we were able to measure C_1/2_ values for 434 protein domains (Fig. 4B). Overall, there was a significant positive correlation between methionines’ PF and C_1/2_ values (Fig. 4C, Spearman Rank Correlation *ρ* = 1.8.10^−10^). As predicted, this correlation was particularly significant for well-buried methionine residues, where most stable protein domains generally contained the most highly protected methionine residues (Fig. 4D). These results demonstrate that thermodynamic folding stabilities can play a major role in establishing the propensity of protein-bound methionines towards oxidation.

### Oxidation protection factors of thermophilic bacteria

In order to verify that thermodynamic folding stabilities can strongly influence methionine oxidation rates, we analyzed the oxidation properties of the highly stable proteome of a model thermophile, *T. thermophilus*. By conducting SPROX experiments, we showed that, as expected, the proteome of *T. thermophilus* has significantly higher C_1/2_ values in comparison to *E. coli* (Fig. 5A). We subsequently quantified methionine PFs at multiple temperatures and observed that the *T. thermophilus* proteome is considerably more resistant to methionine oxidation than the *E. coli* proteome (Fig. 5B). It is important to note that overall, methionines within the *T. thermophilus* and *E. coli* proteomes measured in this study have overall similar SAs (Fig. 5C), *with T. thermophilus* being slightly higher. Hence, the high PF values of the *T. thermophilus* proteome cannot be explained by increased buriedness of methionines and are instead likely due to increased conformational stabilities.

**Figure 5.**
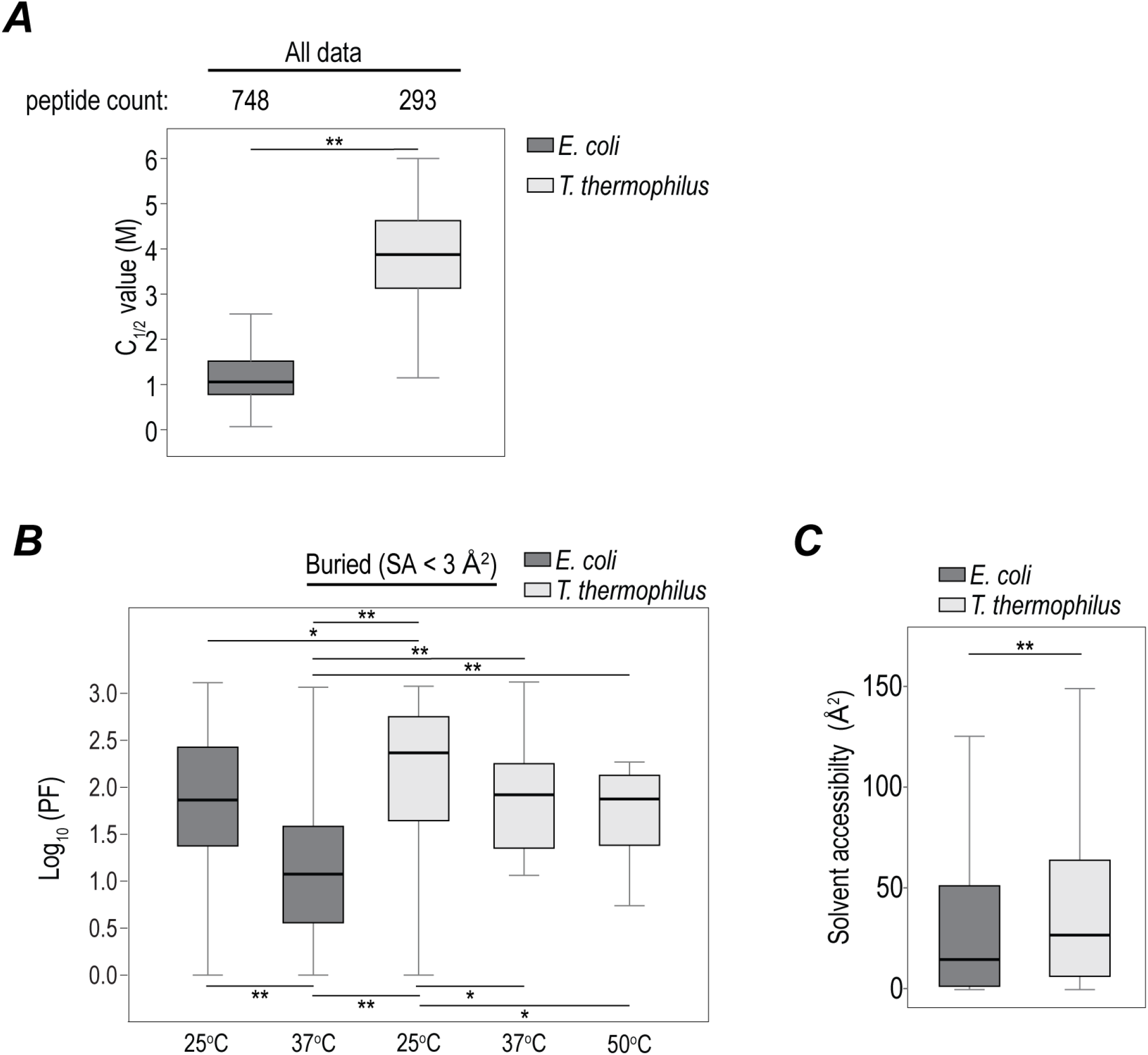
Comparison of C_1/2_ and PF measurements between *E. coli* and *T. thermophilus* proteomes. A) Comparison of the distribution of C_1/2_ values for *E. coli* and *T. thermophilus* at 25°C. B) Comparison of the distributions of PF values for buried methionines within *E. coli* and *T. thermophilus* at varying temperatures. C) Comparison of the distributions of SAs for methionines analyzed in (B). Boxes indicate the interquartile range and whiskers show the entire range of values excluding outliers (> 2 SD). * and ** indicate p-values greater than 0.05 and 0.001, respectively, using a Mann-Whitney U test.

Our observations suggest that oxidation PFs of buried methionines from the native state can be leveraged as a quantitative metric of protein folding stabilities. Considering PFs as a proxy for folding stabilities, our data indicate that of the stabilities of the *E. coli* and *T. thermophilus* proteomes decrease as a function of temperature, and that, at both 25°C and 37°C, the *T. thermophilus* proteome is more stable than the *E. coli* proteome (Fig. 5B). The folding stabilities of the *T. thermophilus* proteome at higher temperatures (37°C and 50°C) are similar to the folding stabilities of the *E. coli* proteome at 25°C. It is interesting to note that *T. thermophilus* is not viable at 25°C, a temperature at which its proteome is highly stable. This observation is consistent with the idea that both destabilization and overstabilization of proteomes may be detrimental to viability. As has been previously suggested, optimal function may require proteomes to retain folded structures, but at marginal stabilities (52).

## Discussion

Our results indicate that protein folding stabilities can play a major role in limiting the oxidation of buried methionine residues. Accordingly, we showed that the proteome of the thermophile *T. thermophilus* is significantly more resistant to oxidation than the mesophile *E. coli*. Interestingly, *T. thermophilus* has the highest levels of oxidation protection at low temperatures where it is unviable. Natural proteins generally have marginal folding stabilities (10 kcal/mol or less). It is generally accepted that the lower limit of protein stability is established by the requirement of folded structures for most protein functions. Additionally, unstable proteins have larger unfolded populations that are nonfunctional and prone to degradation and aggregation. However, reasons for upper limits of protein stability within proteomes are generally less understood and are typically attributed to decreased flexibility that may be required for optimal activity (53,54). Our results suggest an additional explanation as to why over-stabilization may be detrimental to function. Similar to oxidation of buried methionines observed in this study, important PTMs of buried residues may be limited by the folding stability of proteins. Hyper-stable proteomes may be generally less conducive to covalent modifications required for post-translational regulation. This effect may provide one explanation as to why thermophiles are not viable under conditions where their proteomes surpass a maximal stability threshold.

From an application standpoint, our results indicate that oxidation rates of buried methionines from the native state can be used as a convenient proxy for thermodynamic folding stabilities. To date, measurement of protein stabilities using MS-based proteomics has required methodologies that are based on the incremental unfolding of proteins using denaturants or temperature (37,55,56). However, these approaches are complicated by the fact that protein denaturation often results in irreversible aggregation. Furthermore, measurement of protein stability from denaturation curves typically involves making assumptions regarding the two-state nature of protein unfolding and model-dependent extrapolation from measurements under denaturing conditions to native conditions (40,57). Here, we used principles previously outlined for native HDX (50) to show that oxidation rates under native conditions can be used to directly measure protein stabilities without extrapolation from denaturing conditions. This approach should significantly facilitate future proteomic analyses of folding stabilities.

The phenomenon of methionine oxidation is of particular practical interest for the development antibody-based therapeutics where methionine oxidation over long term storage can lead to deactivation and degradation (22,23). Understanding the factors that influence methionine oxidation will enable the identification of labile methionine residues and streamline the development of drug candidates (25,58). The results of this study indicate that enhancing folding stabilities through genetic or chemical approaches may significantly mitigate the detrimental oxidation of protein therapeutics.

## Abbreviations

SA: Solvent-accessible surface area
PF: protection factor
ROS: reactive oxygen species
PTM: post-translational modification
MSR: methionine sulfoxide reductase
MICAL: ‘molecule interacting with CasL’ protein
SPROX: stability of proteins from rates of oxidation

## Acknowledgement

We thank the members of the Ghaemmaghami lab at the University of Rochester for helpful discussions and suggestions.

## Data Availability

All raw and processed data are available in the included Supporting Information and at the ProteomeXchange Consortium via the PRIDE (59) partner repository (accession number PXD030245). Currently, the data can be accessed with the username and password eEcAPYVy.

## Author Information

### Author Contributions

The study concept was conceived by E.W. and S.G. Its detailed planning was performed with contribution from all authors. All wet-lab experiments were conducted by E.W. *T. thermophilus* AlphaFold structures were calculated by A.M.V. and M.O. mass spectrometric analyses were conducted by E.W., K.W., and J.H. Data analysis was conducted by E.W., J.B. and S.G. The manuscript was written by E.W. and S.G. All authors have given approval to the final version of the manuscript.

## Notes

The authors declare no competing financial interest.

## Supporting information

Supporting Table 1 (Supp_table1.xlsx) contains detailed descriptions of proteomic experiments and serves as the key for the raw MS data uploaded to PRIDE.

Supporting Table 2 (Supp_table2.xlsx) contains descriptions and coverage of all MS experiments for each method.

Supporting Table 3 (Supp_table3.xlsx) contains descriptions and lists all of the relevant measured parameters presented in this study that were gathered from our three methods and passed our filtering criteria (see text).

